# Severe introduced predator impacts despite attempted functional eradication

**DOI:** 10.1101/2021.07.19.451788

**Authors:** Brian S. Cheng, Jeffrey Blumenthal, Andrew L. Chang, Jordanna Barley, Matthew C. Ferner, Karina J. Nielsen, Gregory M. Ruiz, Chela J. Zabin

## Abstract

Established non-native species can have significant impacts on native biodiversity without any possibility of complete eradication. In such cases, one management approach is functional eradication, the reduction of introduced species density below levels that cause unacceptable effects on the native community. Functional eradication may be particularly effective for species with limited dispersal ability, which may limit rates of reinvasion from distant populations. Here, we evaluate the potential for functional eradication of introduced predatory oyster drills (*Urosalpinx cinerea*) using a community science approach in San Francisco Bay. We combined observational surveys, targeted removals, and a caging experiment to evaluate the effectiveness of this approach in mitigating the mortality of prey Olympia oysters (*Ostrea lurida*), a conservation and restoration priority species. Despite the efforts of over 300 volunteers that removed over 30,000 oyster drills, we report limited success and discuss several possible mechanisms for this result with broad relevance to management for this and other introduced species. We also found a strong negative relationship between oyster drills and oysters, showing virtually no coexistence across eight sites. At two removal sites, there was no effect of oyster drill removal on oyster survival, which was only observed by caging treatment (0 and 1.6% survival in open and partial cage treatments, as compared to 89.1% in predator exclusion treatments). We conclude that functional eradication of this species requires significantly greater effort and may not be a viable management strategy. Oyster restoration efforts should not be undertaken where *Urosalpinx* is established or is likely to invade.

## INTRODUCTION

Despite widespread efforts to reduce the invasion of non-native species, the rate of introductions appears to be increasing for many taxa (Seebens et al. 2017). The prevention of non-native species introduction is crucial to maintaining and conserving native biodiversity. However, established introduced species can have ongoing negative impacts on native species (Grosholz 2002), and these consequences may be intensified by changing environmental conditions such as climate change (Dukes and Mooney 1999, Sorte et al. 2013). One means of mitigating the impacts of non-native species is managed suppression or “functional eradication” (Hulme 2006, Green et al. 2014), especially when complete eradication may not be achievable. Functional eradication reduces non-native species density below levels that cause unacceptable effects on recipient communities (Green and Grosholz 2021). This approach has been likened to the provisioning of a spatial refuge for native species, such as is the case with protected areas in the context of hunting and extraction (Green et al. 2017).

Functional eradication may be a particularly effective strategy in freshwater and marine habitats where reinvasion rates are high because most taxa produce planktonic dispersive larvae that are cast into the currents (Wray and Raff 1991). The dispersal of these propagules can be several orders of magnitude greater than on land (Kinlan and Gaines 2003), which complicates complete eradication efforts because population rescue can occur via distant populations. However, the potential use of this approach with taxa that have limited dispersal potential and direct development remains less explored. Such species may be good targets for functional eradication, especially when their geographic extent is limited and the probability of reinvasion is low (Liebhold et al. 2016).

In highly invaded San Francisco Bay, the introduced Atlantic oyster drill (*Urosalpinx cinerea;* hereafter *Urosalpinx*) is a direct developing species that is patchily distributed and may be a good candidate for functional eradication for several reasons, including its potential for ecological and economic harm and the potential for success in limiting its abundance and extent. Atlantic oyster drills are predatory snails native to the Atlantic coast of the United States that were introduced to the Pacific and Gulf coasts, as well as parts of Europe beginning in the late 1800s (Fofonoff et al. 2020). *Urosalpinx* is a generalist consumer that can have significant negative impacts on native communities as well as commercial oyster fisheries and aquaculture (Buhle and Ruesink 2009, Kimbro et al. 2009, Koeppel 2011, Cheng and Grosholz 2016). Furthermore, climate change is expected to intensify the impacts of *Urosalpinx* by increasing growth and consumption rates under warming scenarios (Cheng et al. 2017).

*Urosalpinx* has limited dispersal because it lays benthic egg capsules from which fully-formed young emerge (i.e., it does not broadcast spawn its larvae; Carriker 1955). Thus, attempts at removal or managed suppression may not be overwhelmed by reinvasion or recruitment from distant populations (Simberloff 2003). Oyster drills also appear to have limited mobility as adults. In a pilot mark-recapture study of over 500 oyster drills, snails moved a maximum of 4 m from their origin over the course of eight months (A.L. Chang, *unpublished data*). Drills also tend to prefer hard substrate upon which they find prey species and lay egg capsules (Carriker 1955). This can be advantageous in regions where hard substratum is patchy; such is the case in San Francisco Bay, where rocky shores are often isolated by mudflats, which may limit recolonization (immigration) from adjacent habitat patches.

There is also some evidence that key native species may coexist with *Urosalpinx*, including the foundational Olympia oyster (*Ostrea lurida*), a preferred prey species, provided that oyster drill densities remain low (Kimbro and Grosholz 2006, Buhle and Ruesink 2009, Cheng and Grosholz 2016). Bioeconomic models also suggest that the removal of the functionally similar Japanese oyster drill (*Ocinebrellus inornatus*) can be an effective and economical control strategy (Buhle et al. 2005). Moreover, functional eradication may be the only option for controlling introduced species such as *Urosalpinx* in a management landscape with limited resources. Taken together, the biology and management of this system suggests that functional eradication may be a viable option that could have the potential for long lasting and positive effects for native biodiversity.

Community science approaches have garnered much attention as a means of non-native species detection and management (Delaney et al. 2008, Dickinson et al. 2010, Gallo and Waitt 2011). Volunteer-based efforts can vastly increase the spatial and temporal scale of management efforts while reducing financial resources used for control programs (Simberloff 2003). The integration of community members into invasion science also represents an important “social pillar” of sustainable invasive species management (sensu Larson et al. 2011) that can increase social and political capital for such initiatives (Overdevest et al. 2004, Novoa et al. 2018). Community science approaches may also have broader impacts, such as increasing science literacy and the likelihood that participants engage in pro-environmental activities (Crall et al. 2013). At the practical level, community science approaches are a useful management strategy that can have species identification accuracy rates that compare favorably to those of professional scientists (Crall et al. 2011). Volunteer-based hand removal methods can also have high precision and limited environmental impact, as compared to alternatives such as the use of chemical control measures and may be the only feasible approach for mitigating invasive species impacts where such chemicals may not be desirable.

Here, we describe a community science-based approach to attempt the functional eradication of Atlantic oyster drills in Richardson Bay (a sub-embayment of central San Francisco Bay, California USA). San Francisco Bay harbors an overlapping distribution of non-native oyster drills and their prey, the native Olympia oyster. This native oyster is a restoration and conservation priority along the west coast of the U.S. (McGraw 2009, Wasson et al. 2014) that appears to be functionally extinct in many estuaries throughout its range (Zu Ermgassen et al. 2012). San Francisco Bay supports one of the largest remnant Olympia oyster populations throughout its entire range (Polson and Zacherl 2009, Cheng et al. 2016) and invasive oyster drills are thought to be a primary impediment to their restoration and persistence in California (Wasson et al. 2014). Our goals were to (1) quantify the relationship in abundance between non-native oyster drills and native oysters, (2) assess the potential for community science-based removals to reduce oyster drill density, and (3) quantify the effect of removal efforts on oyster survival and growth.

## MATERIALS & METHODS

### Site selection

We focused on Richardson Bay, CA (Fig. 1) because of strong stakeholder interest in the potential long-term observation and maintenance of restoration sites in this region. In addition, the predatory impact of oyster drills had not been quantified anywhere within San Francisco Bay, which supports high native oyster abundance at some sites, primarily where freshwater input may limit drill populations because they are intolerant of low salinity conditions (Cheng et al. 2015, 2017). To broadly quantify the relationship between oyster drills and native oysters, we established eight intertidal field sites. Of these sites, we further focused on a subset of four sites, establishing two sites for community science-based oyster drill removals and two sites as paired controls. One of our removal sites, Lani’s Beach (hereafter Lani’s) was selected because of easy access, high public use, and because it is valuable to the community due to its use in outdoor educational programs at the immediately adjacent Richardson Bay Audubon Center & Sanctuary. If non-native control efforts were successful, this site could be the focus of ongoing community-based invasive species management. Lani’s was paired with a control area that was separated by 50 m along shore (Fig. 1; S1), which greatly exceeds estimated maximum adult movement rates of 4 m over 8 months (A.L. Chang *unpublished data*). Second, we used two sites located on Aramburu Island because of extensive habitat restoration activities that were completed by the Audubon Society in 2010 (Wetlands and Water Resources, Inc. 2010). The Aramburu removal and control sites were separated by ∼140 m along shore (Fig 1, Fig S1). In contrast to Lani’s, Aramburu can only be accessed by boat, making it less frequently visited by the public. Local scale habitat features differ between the sites in that Lani’s is characterized by larger boulders interspersed with mud and cobble, whereas Aramburu is largely characterized by high cover of smaller cobbles with mud located at lower tidal elevations only. Sites also differ in their shoreline orientation which could affect physical characteristics of the environment (e.g., wind exposure, wave energy; Blumenthal 2019).

**Figure 1.**
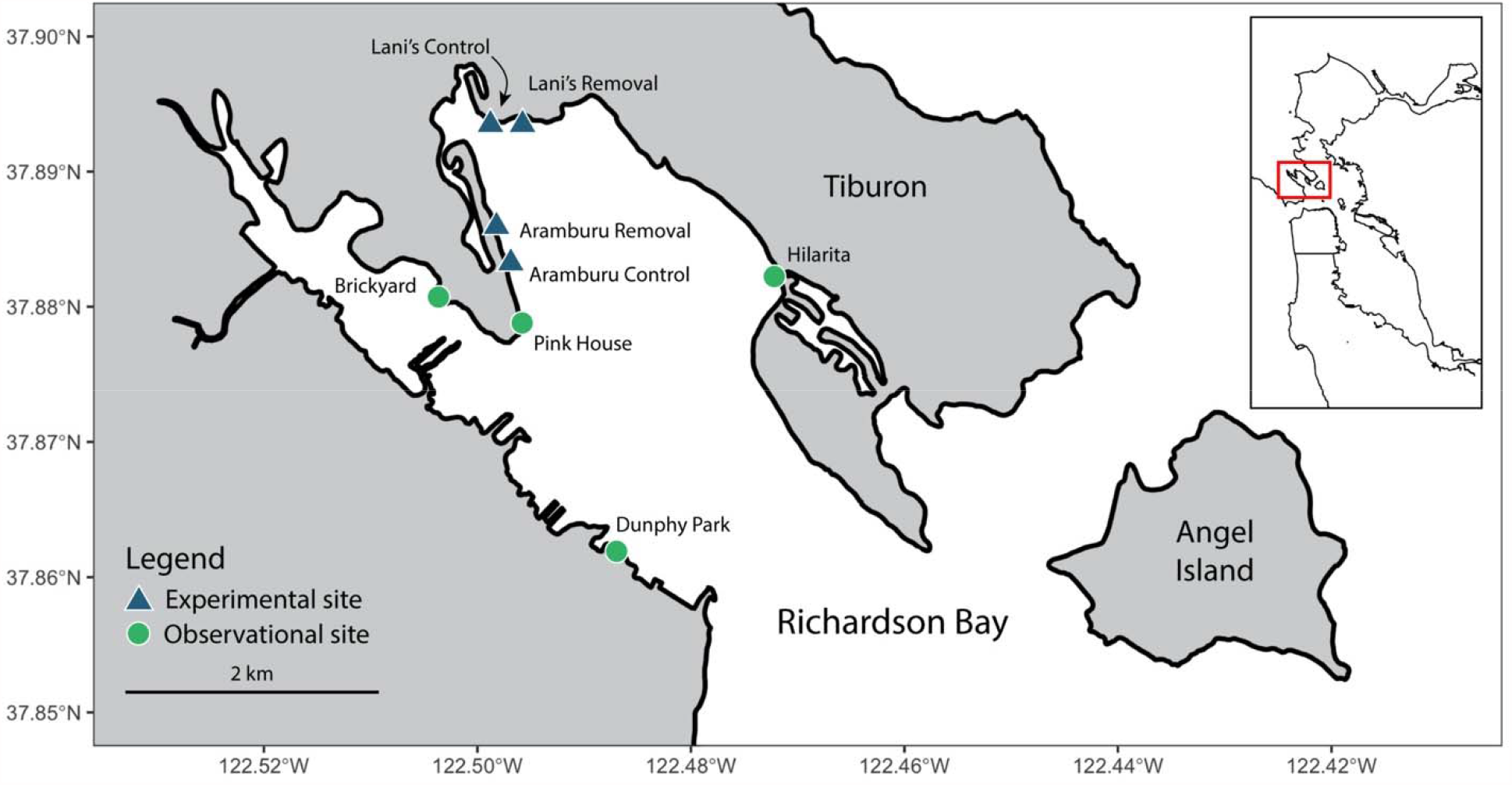
Map of study sites within Richardson Bay, California. Green circles represent sites selected as monitoring locations. Blue triangles represent experimental eradication treatment and control sites. Inset map: the red rectangle depicts location of Richardson Bay within greater San Francisco Bay.

### Observational sampling

To quantify the abundance of *Urosalpinx* and its relationship to native oysters, we conducted quadrat surveys at each of the eight field sites. At each site, we established a permanent 30 m transect at +0.5 m above mean lower low water (MLLW; see Supplementary Information S1 for additional site details). This intertidal elevation typically contains dense native oyster populations in San Francisco Bay and other estuaries (Wasson et al. 2014), and it has hard substrate suitable for oyster recruitment and persistence at our field sites. Along each transect, we censused all oyster drills and oysters at the surface and by overturning all stones within 10 randomly distributed 0.5 ⨯ 0.5 m quadrats. Quadrats were randomly stratified such that five were conducted between 0-15 m of the transect and the remaining five occurred between 15-30 m. For each site, surveys occurred two to five times (20-50 quadrats, mean = 40.5 quadrats) during the low tides of summer, fall, and winter of 2017 and spring and summer of 2018. We then conducted an analysis on a subset of these quadrat data to standardize sampling effort and establish the relationship between drills and oysters prior to any eradication attempts (see Statistics).

### Functional eradication of drills

At each removal site, we established a 60 m swath of shoreline (along shore) from the lower mud zone to the upper barnacle zone (approximately 15 m across shore) to serve as the focal area for removals (see Supplementary Information S1 for maps). For the removal sites, the 60 m total swath included the 30 m fixed transect with additional 15 m buffer zones on each side (along shore). Paired with each eradication site, we established a similar swath of shoreline to serve as a control area except snails were not removed from these areas. Control and removal zones were also separated by stretches of shoreline that did not have hard substrate, potentially limiting the movement of oyster drills. All areas were marked with stakes for the duration of the experiment.

Removals were carried out in the spring with the intent of removing drills before they had laid eggs. We organized drill-removal events, inviting members of the public to assist us in finding and removing drills. To increase community participation, we scheduled drill removals on weekends and only during daylight low tides. In 2017, we organized four removal days at Lani’s and three at Aramburu; in 2018, we held three removal days at each location. Between 20 and 35 people participated at each event and were provided project background and training prior to removal efforts. Teams then worked for 1-2 hours, removing snails by hand. Teams also searched for snails on the surface of the mud, but these were rarely found. To ensure complete spatial coverage of the removal area, we divided removal and buffer areas into ∼2 m wide swaths running perpendicular to shore and assigned volunteers to these zones. For each team, we recorded the number of personnel, their time spent removing drills (h), and the number of snails removed. Collected snails were taken to the Smithsonian Environmental Research Center’s Tiburon, CA laboratory (housed at the Estuary & Ocean Science Center, San Francisco State University) and frozen.

To quantify whether removal efforts were effective at reducing snail densities, we used two approaches. First, we quantified oyster drill densities using quadrats along a fixed transect as in the observational sampling described above. Prior observational data from Tomales Bay suggests that oyster drill and oyster coexistence is possible at an oyster drill density of 5 m^-2^. Experimental data also supports this finding, indicating that the survival of experimentally deployed oyster prey was greater than 50% over 6 months at sites with < 5 drills m^-2^ (Cheng and Grosholz 2016), which we set as our “removal density target”. In 2017, we surveyed before and after each removal event; in 2018 we surveyed immediately before removal events and several days before the oyster outplant experiment described below. In addition to transect surveys, we conducted supplementary surveys at Lani’s where drills aggregate in high densities on several large boulders. Here, we counted drills within 9 haphazardly placed 0.25 m^2^ quadrats placed vertically around the base of each boulder (where drills tended to aggregate). Second, we calculated catch per unit effort (CPUE; number of collected snails per person per hour) from the removal events. We hypothesized that CPUE should decrease over time if functional eradication efforts were successful in depleting the target population.

### Oyster outplant experiment

To determine whether snail removal efforts were sufficient to increase oyster survival, we conducted a field caging experiment in July 2018. This experiment also allowed us to determine whether the physical conditions at each site were able to support oyster survival and growth in the absence of oyster drills. We used the experimental approach from our past efforts to quantify predation intensity in nearby Tomales Bay (Cheng and Grosholz 2016) with a few modifications. First, we constructed experimental units by attaching 8-10 hatchery reared Olympia oysters (Puget Sound Restoration Fund, CA DFW Permit #2018-5211, mean oyster count = 9.8, SD = 0.42) with cyanoacrylate glue to ceramic wall tiles (Daltile model RE1544HD1P4, 10.6 × 10.6 cm). Tiles were individually numbered and held in flow-through seawater tables for two days to verify secure attachment between oysters and tiles. The tiles were also photographed prior to deployment to evaluate potential differences in oyster size, but there was no difference in size across cage treatments (linear mixed model, Wald χ^2^ = 1.1, Df = 3, P = 0.83). We then randomly assigned tiles to one of three treatments: 1) uncaged, fully exposed to predators; 2) caged, no exposure to predators; 3) cage controls. Cages were made of polyethylene aquaculture netting (Memphis Net & Twine PN3, 62.5 mm mesh), wrapped with plastic window screening (Phifer BetterVue Screen, 1 mm mesh), which improved the exclusion of oyster drills in pilot experiments. The cage controls were designed to evaluate cage artifacts, such as shading and reduction of water flow. Cage controls which were identical to the caged treatment except for openings (2.5⨯ 5 cm) that were cut into each cage, which allowed drills access to oysters. Tiles and cages were installed facing horizontally, attached with plastic cable ties to bricks, which were in turn attached to metal rebar driven into the substrate. The bricks were used to keep the experimental units upright and secured to the rebar. Experimental units were randomly stratified by caging treatment type along the +0.5 m MLLW tidal elevation within each removal and control zone at our two field sites (8 replicates × 3 cage treatments × 2 eradication treatments ⨯ 2 sites = 96 experimental units). Tiles and cages were checked one day after deployment (after exposure to two periods of submergence by high tide) to confirm cage integrity.

After 30 days of exposure to field conditions, we recovered 95 of 96 experimental tiles (one uncaged tile was lost) and returned them to the laboratory where each was photographed and examined under a dissecting microscope. Of the 32 caged tiles, we excluded 4 from analysis and graphing because predator exclusion cages were compromised, allowing entry by oyster drills. Oysters were recorded as falling into one of four categories: alive, dead, drilled, or missing. Oysters were scored as alive if the valves retracted upon tapping with a probe. Oysters were classified as dead if no body tissue was found in valves or if the upper valve was removed entirely but the basal valve remained. Oysters were scored as drilled if a bore hole was evident in one of the oyster valves and no body tissue remained. We also quantified oyster growth for surviving oysters from the caged treatment by measuring shell area using image analysis software (ImageJ ver 1.51j8; Schneider et al. 2012). Quantifying oyster growth in the caged treatment allowed us to determine whether the sites were suitable in the absence of oyster drills. Growth was quantified as the final shell area minus the initial shell area for individual surviving oysters.

### Statistics

To quantify the relationship between the abundances of oyster drills and prey oysters, we used a permutation approach to calculate the Spearman rank correlation between their abundances in the quadrat survey data prior to eradication efforts. We used this approach because there was very little coexistence between predator and prey in observational data (i.e. drills and oysters were rarely found in the same quadrat; Fig. 2). This had the effect of generating non-parametric data and many ties in rank that precluded calculation of a p-value for correlation tests. Thus, we randomized (permuted) the data 10,000 times and generated a distribution of Spearman rank correlations and compared it to our original rank correlation to calculate a p-value.

**Figure 2.**
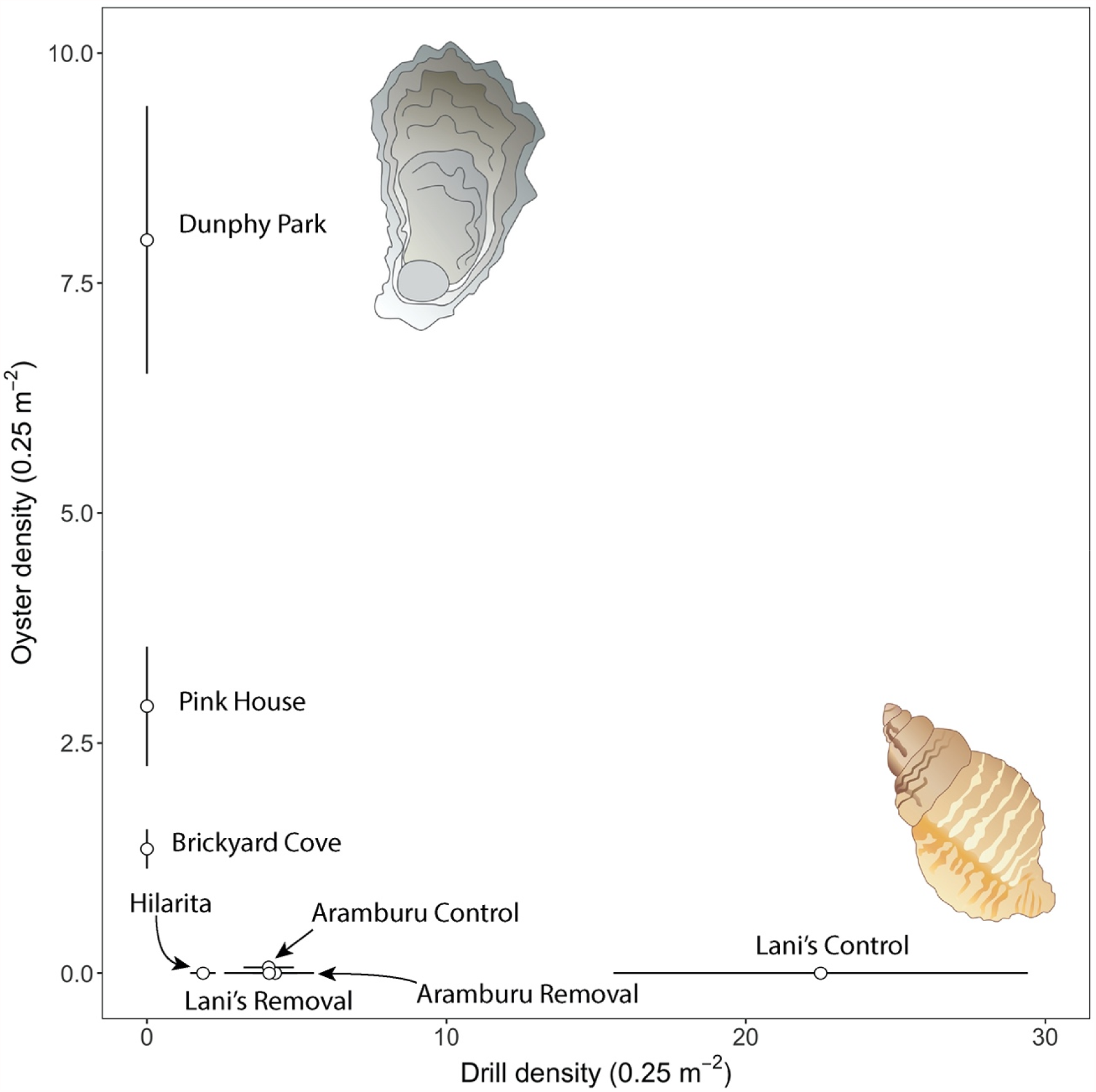
Relationship in abundance between non-native oyster drills and native oysters. There is a strong inverse relationship between the two species where they are rarely found in the same quadrat. Points represent mean counts at each site from quadrat data. Error bars represent standard errors of mean and are calculated for both oysters and drills as double error bars (vertical and horizontal). Most points do not have visible double error bars because of the strong inverse correlation between species. Vector images courtesy of Tracy Saxby and the Integration & Application Network at the University of Maryland Center for Environmental Science.

To evaluate the efficacy of oyster drill eradication efforts, we used two approaches. First, we formulated a removal density target based on prior work from nearby Tomales Bay (Cheng and Grosholz 2016), which demonstrated the potential for oyster drill and oyster coexistence at average drill densities of 5 m^-2^. We then evaluated oyster drill densities at the eradication and control sites with respect to this density removal target using quadrat data after the initiation of removal efforts. Second, we modeled CPUE over time with the expectation that decreased yield would occur once removal efforts began having negative impacts on oyster drill populations. We used generalized linear mixed models to evaluate the fixed effect of time (day of year), site, and their interaction on CPUE (number of drills captured per person). For these models we used a random year effect with a negative binomial error distribution, and an offset term to account for the number of persons and the length of time (person hours) as an index of effort.

To quantify the effect of the removals and cage treatments, we initially used generalized linear mixed models to measure oyster survival. However, the data exhibited complete separation, which occurs when the response data is perfectly predicted by the independent variables (i.e. there is zero variation in a predictor level). Therefore, we used Firth’s bias reduced logistic regression (Heinze and Schemper 2002), which uses a penalized maximum-likelihood estimation procedure. For this analysis, we modeled the effects of experimental manipulation (snail removal vs. control), caging treatment (caged, open, partial), and their interaction as predictors of oyster survival. We modeled surviving and dead oysters at the level of replicate tile (i.e. binomial regression) and assumed that missing oysters were dead. To explore the potential for site-specific differences in environmental conditions to influence oyster performance in the absence of predator effects, we quantified the growth of living oysters within the caged plots (predator exclusion) across all sites. Here we used a linear mixed model with Gaussian error distribution to evaluate the fixed effect of site and the random effect of tile on oyster growth. All analyses and plots were produced in R (ver. 4.0.3; R Core Team 2020), using the packages ‘brglm’, ‘tidyverse’, and ‘glmmTMB’ (Brooks et al. 2017, Wickham et al. 2019, Kosmidis and Firth 2020).

## RESULTS

### Observational sampling

Quadrat surveys revealed a strong inverse relationship between oyster drills and oysters across the eight monitoring sites (Fig. 2). At 7 of 8 sites, quadrats revealed the presence of oysters or oyster drills but never both. In fact, only 2 out of 324 quadrats contained at least one live oyster and one live drill (both quadrats at Aramburu control). The randomization test revealed strong evidence for a negative relationship between oysters and drills (ρ = -0.55, P <0.001).

### Functional eradication of drills

Over 6 events in 2017 and 2018 at the Aramburu removal site, we recruited 115 participants who worked 183 hours to remove 12,261 oyster drills. Over 7 events in 2017 and 2018 at Lani’s, we recruited 202 participants who worked 284 hours to remove 19,297 oyster drills. Success in achieving the density removal target was limited. The target of 5 oyster drills m^-2^ was only achieved at 1 of 6 possible time points (excluding the survey prior to removal efforts) at Aramburu (Fig. 3). At Lani’s the target removal density was achieved at 3 of 7 possible time points in cobble habitat and at 0 of 7 time points in boulder habitat (Fig. 3). In addition, drill densities were well above the target density prior to the July 2018 oyster outplant experiment, averaging 16.4 drills m^-2^ at Aramburu and 15.2 and 207.1 drills m^-2^ at Lani’s cobble and boulder habitat, respectively. There was no evidence that oyster drill CPUE changed over time at either site (Table 1, Fig. 4).

**Table 1.** Statistical results from multiple surveys and experiments. A) Catch per unit effort modeling, testing for a change in CPUE over time (day of year) across sites. CPUE is modeled as number of drills caught standardized by effort with a person-hour offset. B) Results from bias reduced logistic regression for the oyster caging experiment. The caged plots at control sites are the intercept and all parameter estimates are in relation to this group. The model estimates survival, so a negative estimate indicates that survival declines in relation to the caged treatment (i.e. for open and partial cage treatments). Only caged treatments had an effect on oyster survival, whereas oyster drill eradication had no effect.

| Parameters | Estimate | SE | z-value | P |
| --- | --- | --- | --- | --- |
| **A) Catch per unit effort** |  |  |  |  |
| Aramburu (intercept) | 4.17 | 0.49 | 8.5 | <0.001 |
| Day of year | 0.00 | 0.01 | 0.12 | 0.90 |
| Lani's | 0.62 | 0.57 | 1.11 | 0.27 |
| Day of year * Lani's | -0.01 | 0.01 | -0.89 | 0.38 |
| **B) Oyster caging experiment** |  |  |  |  |
| Caged (intercept) | 2.14 | 0.30 | 7.14 | <0.001 |
| Open | -7.84 | 1.45 | -5.40 | <0.001 |
| Partial cage | -5.67 | 0.57 | -10.02 | <0.001 |
| Eradication | -0.11 | 0.39 | -0.30 | 0.77 |
| Open x Eradication | 0.07 | 2.05 | 0.03 | 0.97 |
| Partial cage x Eradication | -1.01 | 1.03 | -0.98 | 0.33 |

**Figure 3.**
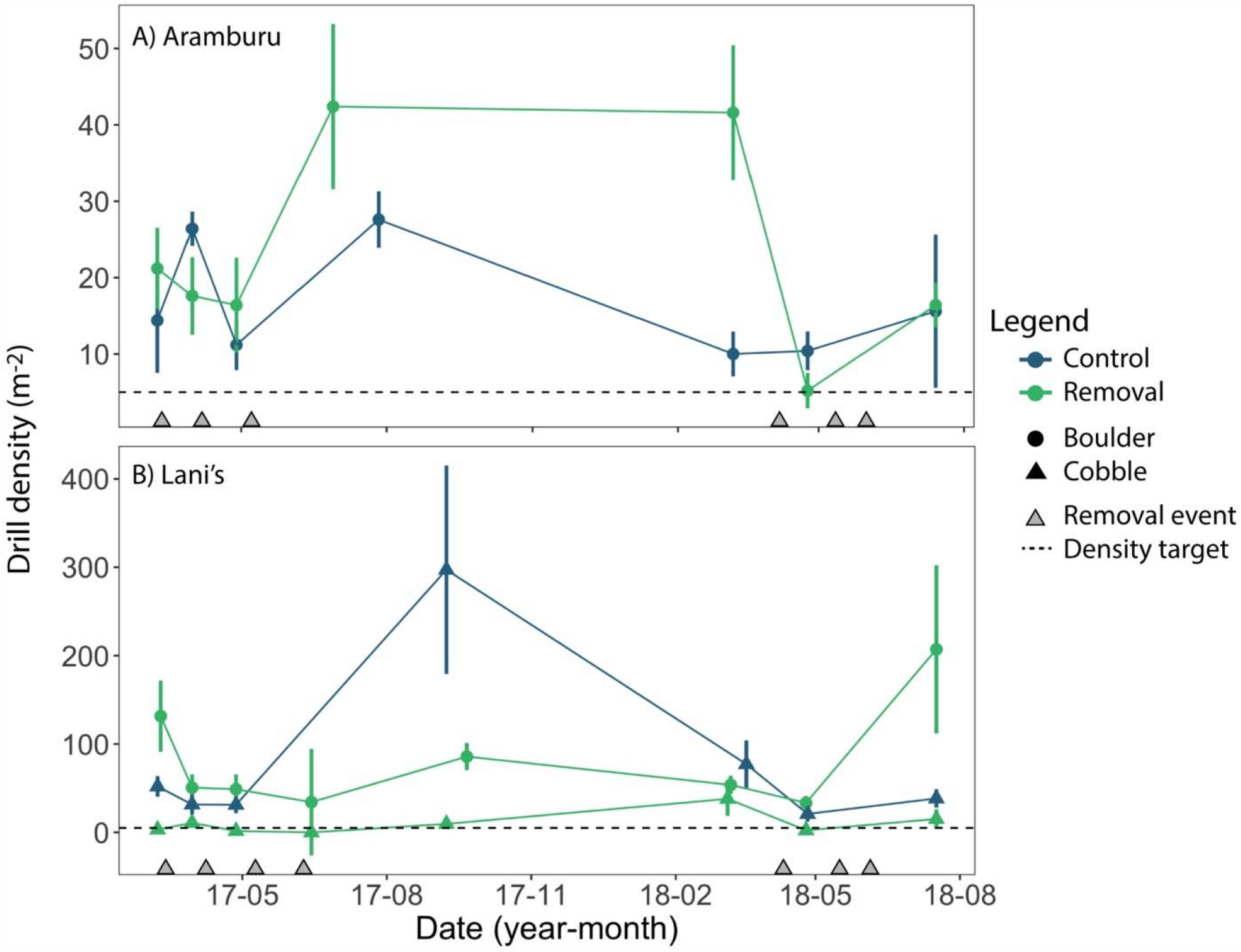
Oyster drill densities across control and removal sites at A) Aramburu and B) Lani’s. Point and error estimates refer to mean ± SEM. Dashed line represents the density target based on coexistence of oyster drills and oysters in nearby Tomales Bay. Note the different y-axis scales on each panel.

**Figure 4.**
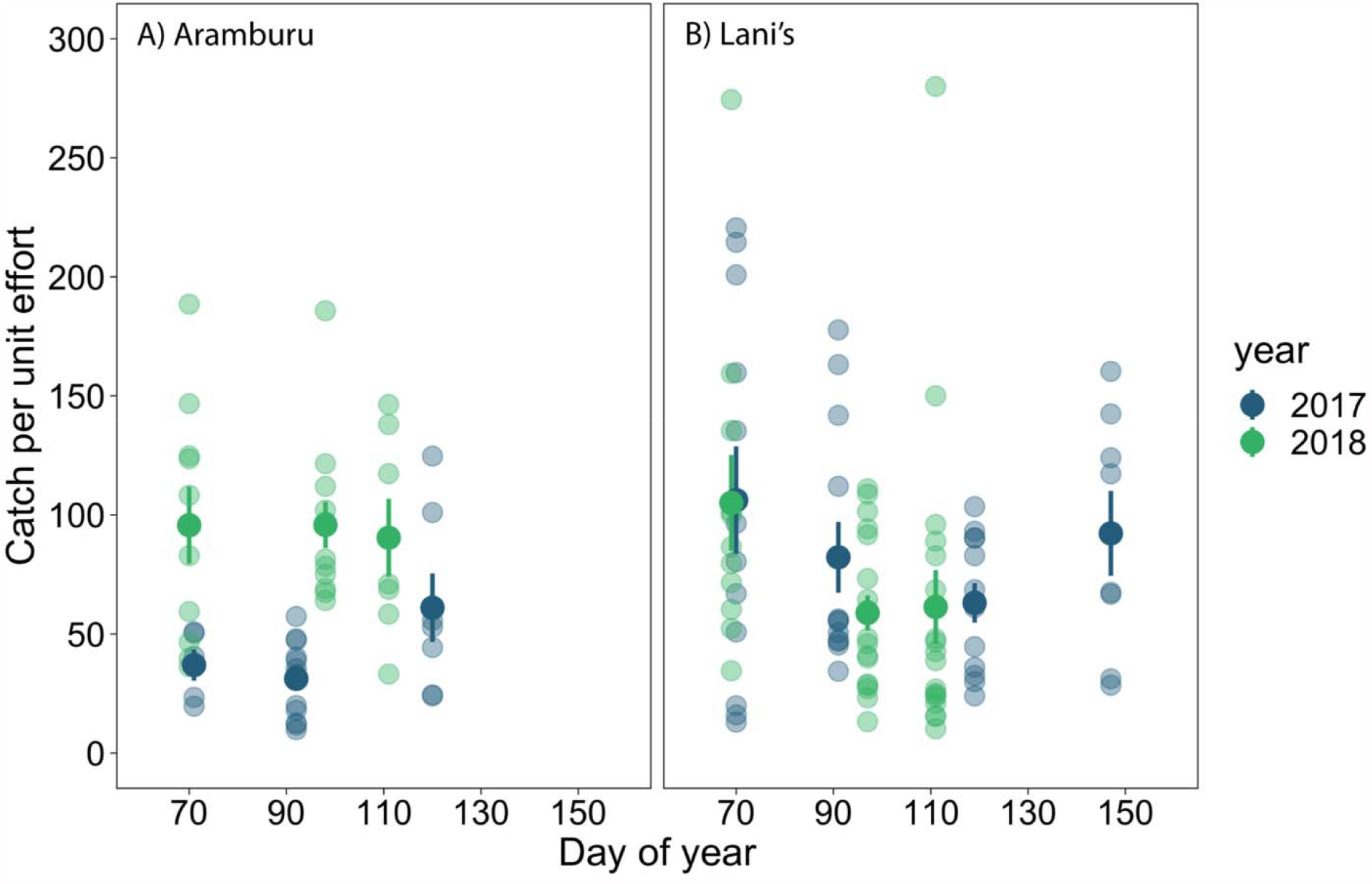
Catch per unit effort (CPUE) across day of year at each eradication site in 2017 and 2018. CPUE is calculated as number of drills captured per person-hour. Each point refers to a CPUE estimate for individual volunteer groups. Bolded point and error bar refers to mean CPUE and standard error of the mean for that eradication day. We hypothesized that CPUE would show a downward trend over time if removals were causing declines in population abundance.

### Oyster outplant experiment

Oyster drill removal efforts did not appear to affect oyster survival. Survival in the oyster outplant experiment was highly dependent on caging treatment but not the removal treatment, nor their interaction (Table 1, Fig. 5-6). In the caged plots, overall survival was 89.1% (246 of 276 oysters), whereas survival was 0% (0 of 305 oysters) and 1.6% (5 of 315 oysters) in the open and partial caged plots, respectively (Fig. 6). The number of drilled oysters also differed based on cage treatment. We found 0 drilled oysters in caged plots, as opposed to 148 (48.5%) and 127 (40.3%) in open and cage control plots, respectively (Supplementary Information S2). Oysters from the caged plots not only survived well across sites but also exhibited high growth that ranged from 1.8-2.3 cm^2^ (378-456%) over the 30-day experiment (Fig. 7). There was some marginal evidence for differences in growth across sites (Wald χ^2^ = 6.9, df = 3, P = 0.075) but differences were small and all locations supported positive and high levels of growth.

**Figure 5.**
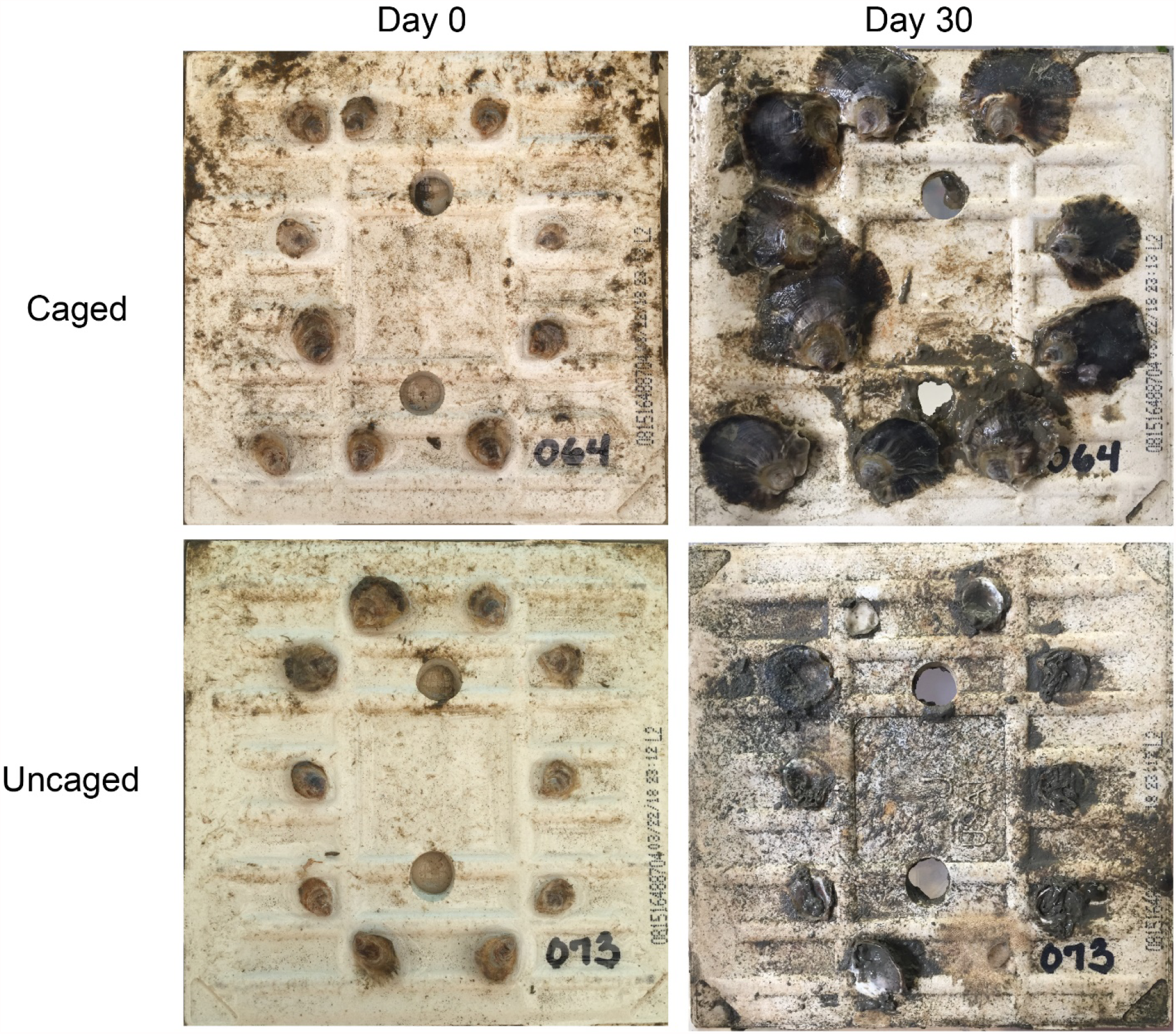
Sample photographs of oyster tiles before (day 0) and after field deployment (day 30). The caged tile was protected from predators whereas the uncaged tile was exposed. The center bore holes were used to affix the tile to the cage and/or rebar stake.

**Figure 6.**
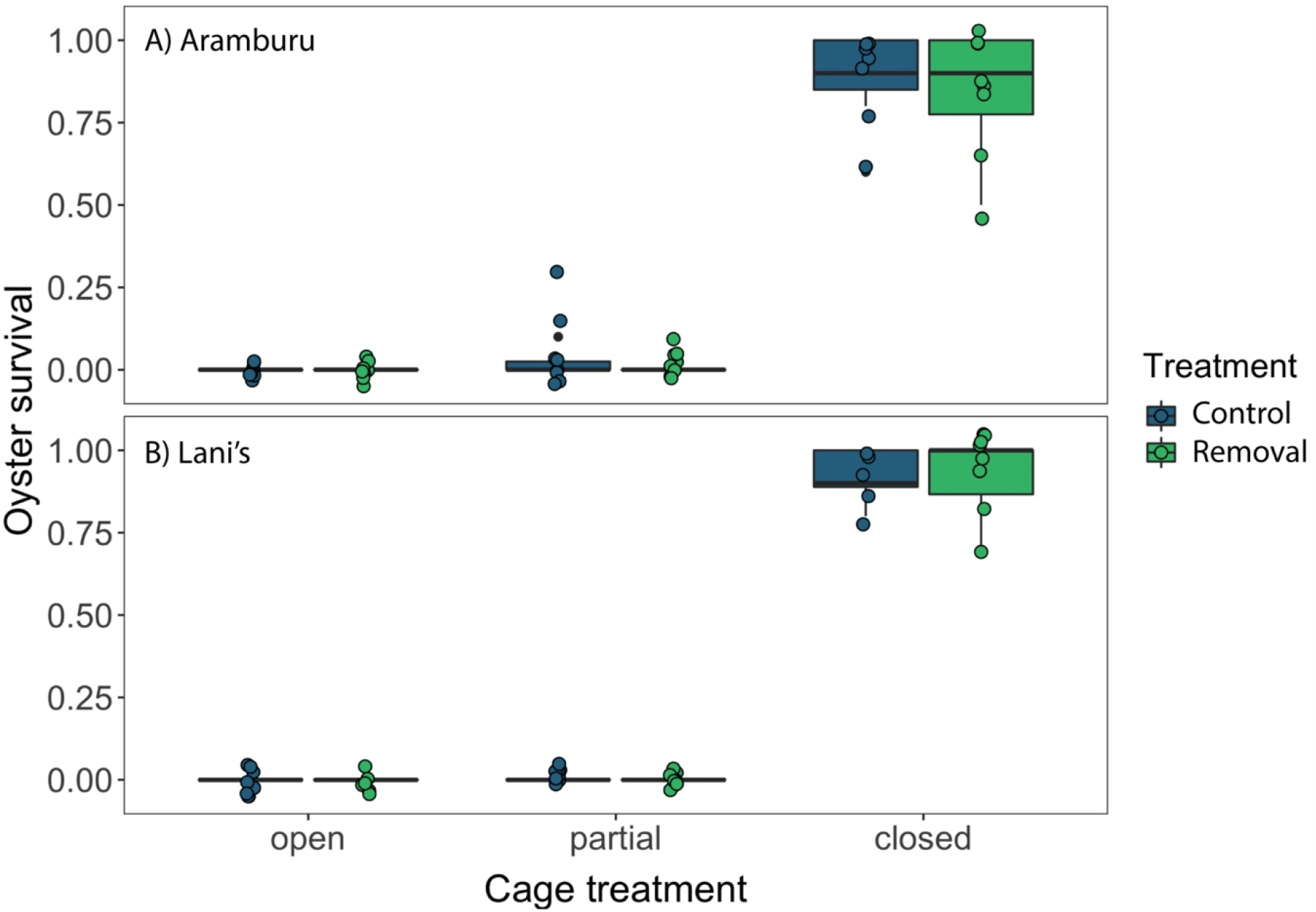
Oyster survival across caging treatments at A) Aramburu and B) Lani’s at control and removal sites. Survival is plotted with boxplots and overlaid jittered points. Proportional survival was calculated at the tile level (N = 5-8 tiles per treatment). Boxplots collapse to a solid line because of zero variation (and zero survival) in most open and partial cage plots.

**Figure 7.**
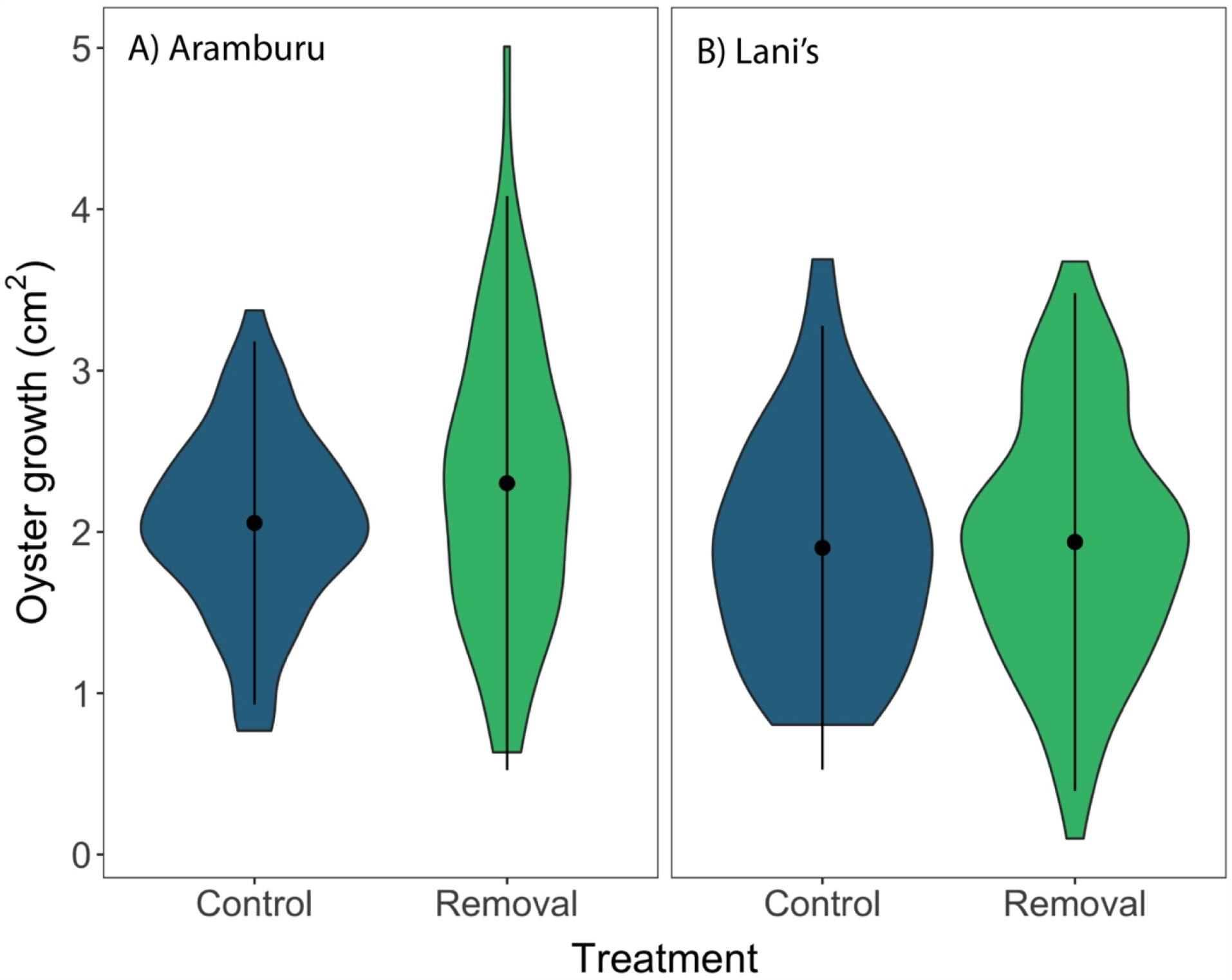
Violin plots of oyster growth from caged (predator exclusion) tiles across sites at A) Aramburu and B) Lani’s. Point and error estimates within each violin refer to the mean ±SD. N = 68 and 67 oysters at Aramburu control and removal, respectively. N = 43 and 74 oysters at Lani’s control and removal, respectively.

## DISCUSSION

Even though oyster drills have life history traits that appear ideal for functional eradication, our efforts to manage this species resulted in limited success. Removals of over 30,000 oyster drills from Aramburu and Lani’s partly reduced oyster drill abundances but were insufficient to reach target oyster drill densities, only achieving this goal for 4 of 13 total survey time points in cobble habitat. We also saw no evidence for declining CPUE, which would have been expected if oyster drill populations were undergoing significant depletion. Consequently, the survival of Olympia oysters in our outplant experiment was extremely low and unaffected by the removal treatment. In contrast, survival was highly dependent on the caging treatment (high survival only in predator exclusion treatments) where up to 48.5% of oysters had drill holes, highlighting the role of predator-induced mortality by drills. There was also no evidence that environmental site-specific differences could have driven oyster mortality. Oysters in the predator exclusion treatment exhibited high survivorship and high growth across all sites. Taken together with our observational surveys which revealed a strong negative relationship between oysters and predatory oyster drills, these data suggest large negative effects of non-native *Urosalpinx* on Olympia oysters in Richardson Bay and likely other regions of San Francisco Bay (Wasson et al. 2014).

Future functional eradication efforts are likely to be most successful if the mechanisms underlying population persistence of the introduced species are well understood. Efforts to control other introduced marine species have been successful at low population sizes and early stages of invasion. For example, successful control efforts for non-native mussels in Spain used community science approaches to remove approximately 800 mussels in one event at 1-2 years after initial detection (Miralles et al. 2016). Likewise, community science-based control efforts with lionfish “derbies” have been successful in areas with annual removals on the order of 1,000 fish (Green et al. 2017). By contrast, our efforts over 3-4 removal events yielded 5,000 -10,000 oyster drills per year at each site but without lasting impact on local densities, suggesting a much greater total population size for these established *Urosalpinx* populations. Such results highlight the importance of focused high-intensity control efforts soon after introduction when abundance and population persistence may be low (Simberloff et al. 2013). The above discussed removal efforts may also have been aided by the lack of a temporal or spatial refuge (e.g., sessile or tropical species). In contrast, our removal efforts may have been further compounded by the overwintering of oyster drills. During colder winter months, drills reduce their activity and may burrow within sediments which could provide a spatial and temporal refuge from removal efforts not seen with sessile species such as mussels (Carriker 1955). We aimed for spring removals, prior to initiation of egg clutch laying to prevent the recruitment of young drills, but it was also possible that not all drills were active in earlier removal events, reducing the efficacy of these efforts. Another possible mechanism limiting our success may have been reinvasion of oyster drills from neighboring habitat via emigration, as opposed to larval dispersal. This could have occurred if drills were located at intertidal heights below our focal removal area or adjacent to our focal removal areas in the alongshore direction. Given the limited movement of oyster drills observed in pilot mark-recapture experiments, this mechanism would seem possible but less likely. We also observed an initial decrease in oyster drill density at Aramburu concomitant with the removal events, followed by an increase (Fig. 3A). The mechanism for this pattern is unclear but could have been linked to “overcompensation” or the “hydra effect”, which occurs when removals counterintuitively result in population increases (Roos et al. 2007, Abrams 2009). Such a phenomenon can arise if removal reduces the strength of negative intraspecific interactions (e.g. competition or cannibalism) driving greater population growth, as seen in fish harvesting efforts in lakes (Zipkin et al. 2008) and the removal of invasive European green crabs (Grosholz et al. 2021).

Instead of functional eradication, greater success with oyster drill control might be achieved via species interactions with native species (i.e., biotic resistance; Kimbro et al. 2009). Native rock crabs (*Romaleon antennarius* and *Cancer productus*) are generalist consumers that prey upon oyster drills in laboratory and field experiments and appear to set the local range limit of oyster drills (*Urosalpinx* and *Ocinebrellus inornatus*) in Tomales Bay (Cheng and Grosholz 2016). Although we did not quantify native crab densities in this study, we rarely observed these crab species at our sites and observed extremely high oyster drill densities as compared to other estuaries, suggesting that oyster drills experience low predator-induced mortality. Given the coastal distribution of these crabs and evidence for lower thermal preferences (Sulkin and McKeen 1994, Padilla-Ramírez et al. 2015), one possibility is that rock crabs are thermally limited in Richardson Bay. These sites are located adjacent to broad intertidal mudflats which may limit the movements of crabs and the accessibility of thermal refugia (e.g. deep, cooler waters). In addition, crabs themselves may be subject to predator-induced mortality from birds, sharks, and rays (Gray et al. 1997, Ebert and Ebert 2005). Future oyster restoration efforts may benefit from a greater mechanistic understanding of the forces limiting crab abundance to take advantage of the trophic cascades that these predators can generate (Kimbro et al. 2009, Cheng and Grosholz 2016).

Oyster drills have widespread predatory effects on native oysters throughout many estuaries in California, including Tomales Bay (Kimbro et al. 2009, Cheng and Grosholz 2016), Humboldt Bay (Koeppel 2011), and San Francisco Bay (this study). Oyster drills have are also established in Willapa Bay, WA although they appear to have a smaller role in driving oyster mortality there (Buhle and Ruesink 2009). *Urosalpinx* impacts include lethal, as well as non-lethal effects, such as induced shell thickening in the presence of oyster drills (Bible et al. 2017). Evidence from this study and others suggests oyster restoration at sites with established populations of oyster drills are unlikely to be successful. Oyster populations at central San Francisco Bay sites, which experience more frequent low salinity periods, appear to have refuge from predators because the salinity tolerance of drills is lower than that of oysters (Cheng et al. 2017). However, such sites can also experience both extreme low-salinity events that can drive oyster mass-mortality events (Cheng et al. 2016) and extreme variation in oyster recruitment (Chang et al. 2018). Alternatively, lower estuary sites may be suitable if they can support crab populations that can suppress oyster drill abundance. However, these sites may have diminished recruitment of oyster larvae driven by greater coastal influence and reduced residence time of water masses (Wasson et al. 2016, Chang et al. 2018). Given the dynamic variation of physical and abiotic conditions within estuaries, the impacts of oyster drills, and therefore success of oyster restoration efforts will be highly dependent on within estuary siting.

One largely successful aspect of this project was the engagement with community members. This project recruited over 300 volunteers in our outreach and research activities, highlighting the significant interest in non-native species control and shoreline restoration, and a small group of volunteers enthusiastically continued to remove snails from Lani’s, removing another ∼13,000 snails in 2019. Such a partnership is critical for building science knowledge among the public and garnering public support for environmental science issues (Larson et al. 2011, Novoa et al. 2018). Community science partnerships have grown considerably in the last several decades, and this project serves as a regional model for interacting with the community in a science-based context.

Overall, our study highlights the difficulties of managing established and highly abundant introduced species that have negative effects on native communities. For well-established and successful species, functional eradication may be a viable option for suppressing introduced species, but there should be careful consideration of the species’ life history and environmental conditions in determining project success. In situations where functional eradication may not be feasible, promoting biotic resistance by native enemies and focusing on containment may be more desirable uses of the limited time and resources available to manage highly successful invaders whose impacts may intensify under radically changing environmental conditions.

## ACKNOWLEDGEMENTS

We thank all the community volunteers for their efforts in removing oyster drills from the study sites. We also thank the Marin Community Foundation and Buck Family Foundation for granting funds administered through the California State Coastal Conservancy’s Advancing Nature-Based Adaptation Solutions in Marin County program. This project was also supported by a gift from Jason Payne. We also thank our collaborators at the Richardson Bay Audubon Center, including Courtney Gutman. We thank Alison Cawood, Jason Thomas, and Evie Borchard for assistance in the field and organizing volunteer efforts. Support for M.C.F was provided by an award under the Federal Coastal Zone Management Act, administered by the National Oceanic and Atmospheric Administration’s Office for Coastal Management (to San Francisco State University).

---

**Supplementary Information S1.**
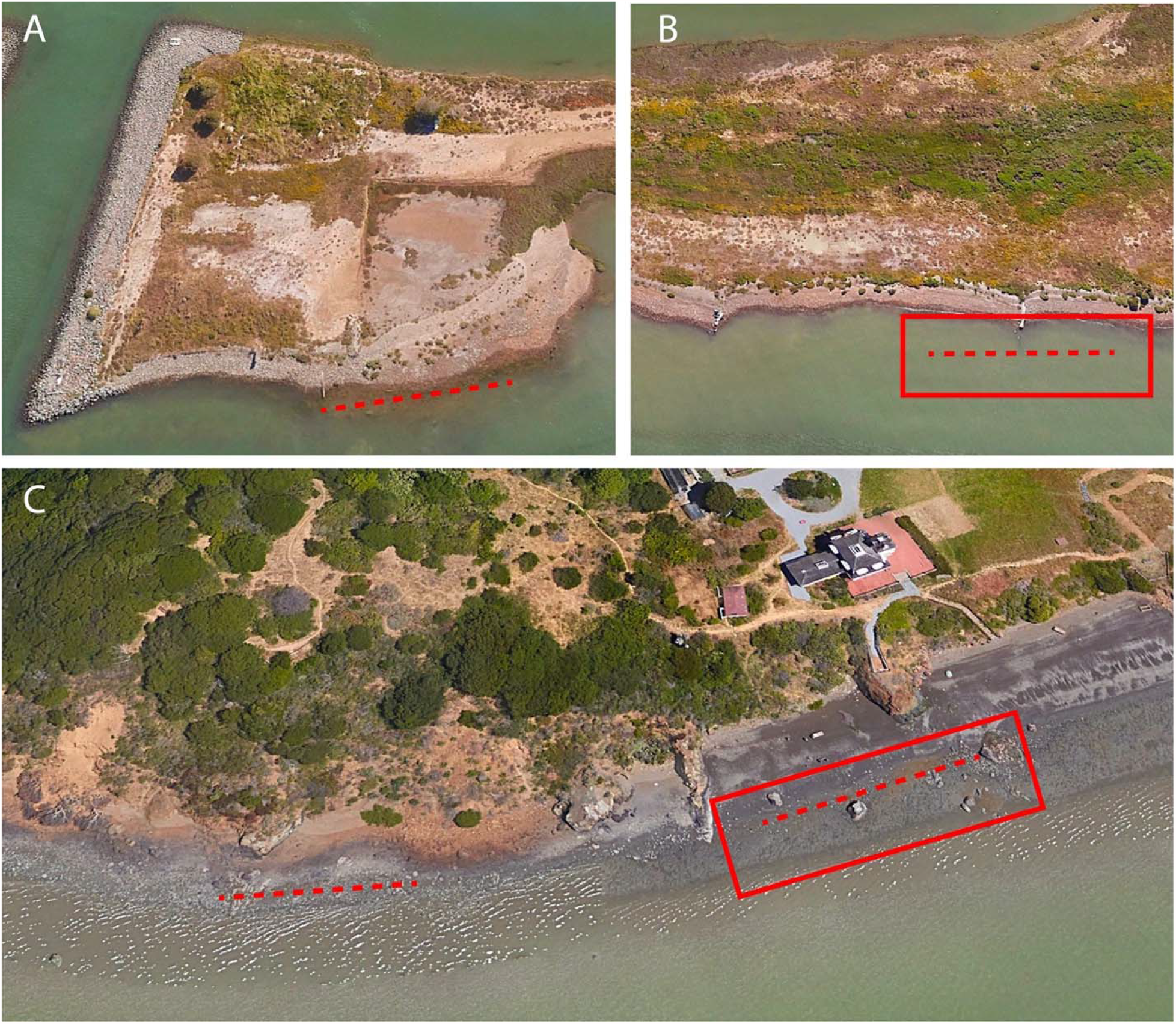
Specific location of observational transects (dashed lines, 30 m length) and eradication zones (solid rectangles, 60 × 15 m) at **A)** Aramburu Control, **B)** Aramburu Removal, and **C)** Lani’s Control and Removal. Lines and rectangles are approximately drawn. *Google Earth*, earth.google.com/web (June 10, 2019). Eye altitude 550 feet.

**Supplementary Information S2.**
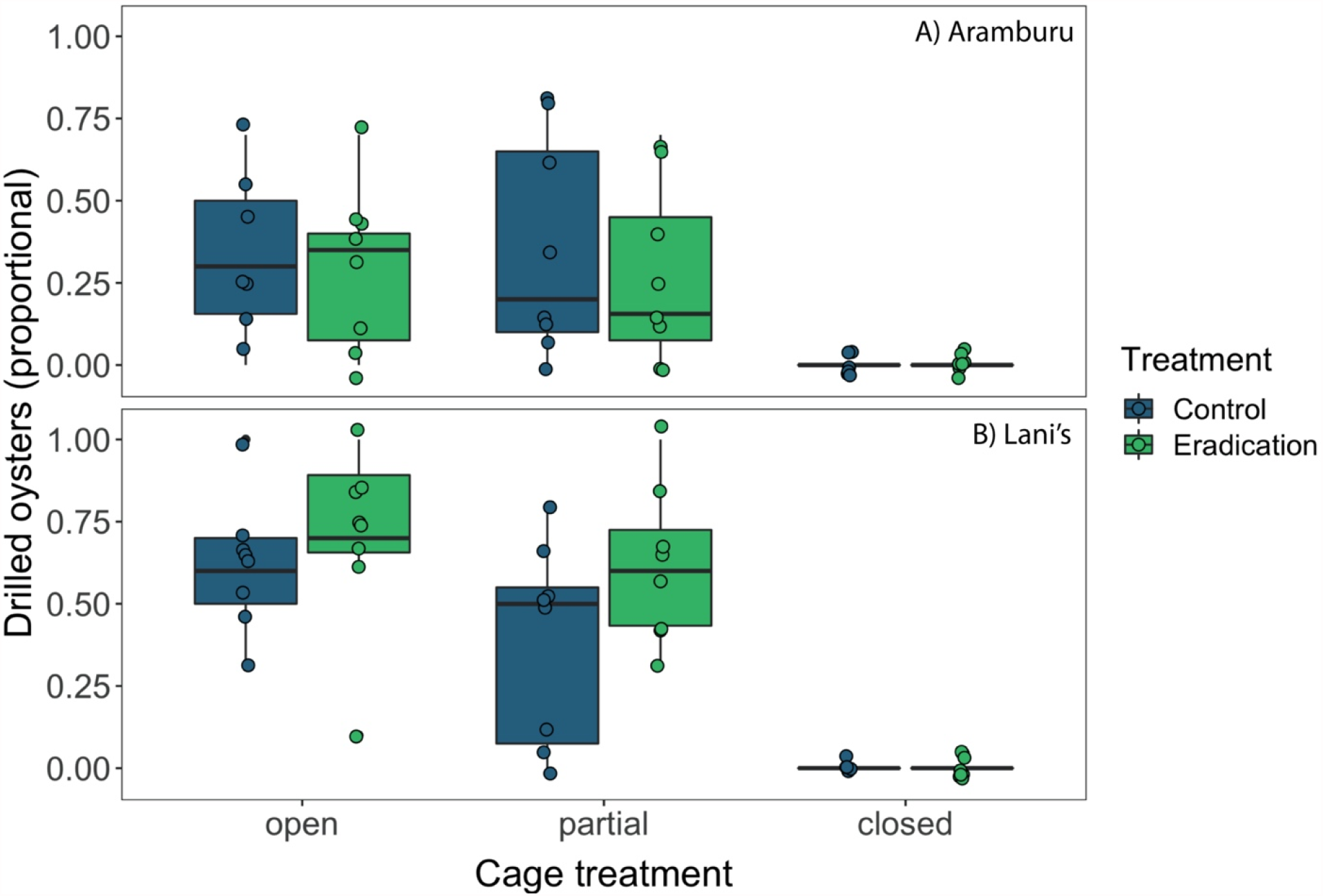
Drilled oysters across caging treatments at A) Aramburu and B) Lani’s at control and removal sites. Drilled oysters are plotted with boxplots and overlaid jittered points. Drilled oysters were calculated as a proportion (number of oysters possessing at least one drill hole divided by the initial number of oysters at the tile level). N = 5-8 tiles per treatment. Boxplots collapse to a solid line because of zero variation (and zero survival) in all closed plots.

## REFERENCES

Abrams, P. A. 2009. When does greater mortality increase population size? The long history and diverse mechanisms underlying the hydra effect. Ecology Letters 12:462–474.

Bible, J. M., K. R. Griffith, and E. Sanford. 2017. Inducible defenses in Olympia oysters in response to an invasive predator. Oecologia 183:809–819.

Blumenthal, J. 2019. Modeling habitat covariates for Atlantic oyster drills in Richardson Bay, California. Master’s Thesis, San Francisco State University.

Brooks, M. E., K. Kristensen, K. J. Van Benthem, A. Magnusson, C. W. Berg, A. Nielsen, H. J. Skaug, M. Machler, and B. M. Bolker. 2017. glmmTMB balances speed and flexibility among packages for Zero-inflated Generalized Linear Mixed Modeling. The R Journal 9:378–400.

Buhle, E. R., M. Margolis, and J. L. Ruesink. 2005. Bang for buck: cost-effective control of invasive species with different life histories. Ecological Economics 52:355–366.

Buhle, E. R., and J. L. Ruesink. 2009. Impacts of invasive oyster drills on Olympia oyster (Ostrea lurida Carpenter 1864) recovery in Willapa Bay, Washington, United States. Journal of Shellfish Research 28:87–96.

Carriker, M. R. 1955. Critical review of biology and control of oyster drills Urosalpinx and Eupleura. U.S. Fish and Wildlife Service Special Scientific Report 148:1–151.

Chang, A. L., A. K. Deck, L. J. Sullivan, S. G. Morgan, and M. C. Ferner. 2018. Upstream— Downstream Shifts in Peak Recruitment of the Native Olympia Oyster in San Francisco Bay During Wet and Dry Years. Estuaries and Coasts 41:65–78.

Cheng, B. S., J. M. Bible, A. L. Chang, M. C. Ferner, K. Wasson, C. J. Zabin, M. Latta, A. Deck, A. E. Todgham, and E. D. Grosholz. 2015. Testing local and global stressor impacts on a coastal foundation species using an ecologically realistic framework. Global Change Biology 21:2488–2499.

Cheng, B. S., A. L. Chang, A. Deck, and M. C. Ferner. 2016. Atmospheric rivers and the mass mortality of wild oysters: insight into an extreme future? Proceedings of the Royal Society of London B: Biological Sciences 283:20161462.

Cheng, B. S., and E. D. Grosholz. 2016. Environmental stress mediates trophic cascade strength and resistance to invasion. Ecosphere 7:e01247.

Cheng, B. S., L. M. Komoroske, and E. D. Grosholz. 2017. Trophic sensitivity of invasive predator and native prey interactions: integrating environmental context and climate change. Functional Ecology 31:642–652.

Crall, A. W., R. Jordan, K. Holfelder, G. J. Newman, J. Graham, and D. M. Waller. 2013. The impacts of an invasive species citizen science training program on participant attitudes, behavior, and science literacy. Public Understanding of Science 22:745–764.

Crall, A. W., G. J. Newman, T. J. Stohlgren, K. A. Holfelder, J. Graham, and D. M. Waller. 2011. Assessing citizen science data quality: an invasive species case study. Conservation Letters 4:433–442.

Delaney, D. G., C. D. Sperling, C. S. Adams, and B. Leung. 2008. Marine invasive species: validation of citizen science and implications for national monitoring networks. Biological Invasions 10:117–128.

Dickinson, J. L., B. Zuckerberg, and D. N. Bonter. 2010. Citizen Science as an Ecological Research Tool: Challenges and Benefits. Annual Review of Ecology, Evolution, and Systematics 41:149–172.

Dukes, J. S., and H. A. Mooney. 1999. Does global change increase the success of biological invaders? Trends in Ecology & Evolution 14:135–139.

Ebert, D. A., and T. B. Ebert. 2005. Reproduction, diet and habitat use of leopard sharks, Triakis semifasciata (Girard), in Humboldt Bay, California, USA. Marine and Freshwater Research 56:1089–1098.

Encarnação, J., M. A. Teodósio, and P. Morais. 2021. Citizen Science and Biological Invasions: A Review. Frontiers in Environmental Science 8.

Gallo, T., and D. Waitt. 2011. Creating a Successful Citizen Science Model to Detect and Report Invasive Species. BioScience 61:459–465.

Gray, A. E., T. J. Mulligan, and R. W. Hannah. 1997. Food habits, occurrence, and population structure of the bat ray, Myliobatis californica, in Humboldt Bay, California. Environmental Biology of Fishes 49:227–238.

Green, S. J., N. K. Dulvy, A. M. L. Brooks, J. L. Akins, A. B. Cooper, S. Miller, and I.M. Côté. 2014. Linking removal targets to the ecological effects of invaders: a predictive model and field test. Ecological Applications 24:1311–1322.

Green, S. J., and E. D. Grosholz. 2021. Functional eradication as a framework for invasive species control. Frontiers in Ecology and the Environment 19:98–107.

Green, S. J., E. B. Underwood, and J. L. Akins. 2017. Mobilizing volunteers to sustain local suppression of a global marine invasion. Conservation Letters 10:726–735.

Grosholz, E. 2002. Ecological and evolutionary consequences of coastal invasions. Trends in Ecology & Evolution 17:22–27.

Grosholz, E., G. Ashton, M. Bradley, C. Brown, L. Ceballos-Osuna, A. Chang, C. de Rivera, J. Gonzalez, M. Heineke, M. Marraffini, L. McCann, E. Pollard, I. Pritchard, G. Ruiz, B. Turner, and C. Tepolt. 2021. Stage-specific overcompensation, the hydra effect, and the failure to eradicate an invasive predator. Proceedings of the National Academy of Sciences 118.

Heinze, G., and M. Schemper. 2002. A solution to the problem of separation in logistic regression. Statistics in Medicine 21:2409–2419.

Hulme, P. E. 2006. Beyond control: wider implications for the management of biological invasions. Journal of Applied Ecology 43:835–847.

Kimbro, D. L., and E. D. Grosholz. 2006. Disturbance influences oyster community richness and evenness, but not diversity. Ecology 87:2378–2388.

Kimbro, D. L., E. D. Grosholz, A. J. Baukus, N. J. Nesbitt, N. M. Travis, S. Attoe, and C. Coleman-Hulbert. 2009. Invasive species cause large-scale loss of native California oyster habitat by disrupting trophic cascades. Oecologia 160:563–575.

Kinlan, B. P., and S. D. Gaines. 2003. Propagule Dispersal in Marine and Terrestrial Environments: A Community Perspective. Ecology 84:2007–2020.

Koeppel, J. A. 2011. High Predation May Hinder Native Oyster (Ostrea lurida Carpenter, 1864) Restoration in North Humboldt Bay, California. PhD Thesis, Humboldt State University.

Kosmidis, I., and D. Firth. 2020. Jeffreys-prior penalty, finiteness and shrinkage in binomial-response generalized linear models. Biometrika:asaa052.

Larson, D. L., L. Phillips-Mao, G. Quiram, L. Sharpe, R. Stark, S. Sugita, and A. Weiler. 2011. A framework for sustainable invasive species management: Environmental, social, and economic objectives. Journal of Environmental Management 92:14–22.

Liebhold, A. M., L. Berec, E. G. Brockerhoff, R. S. Epanchin-Niell, A. Hastings, D. A. Herms, J. M. Kean, D. G. McCullough, D. M. Suckling, P. C. Tobin, and T. Yamanaka. 2016. Eradication of Invading Insect Populations: From Concepts to Applications. Annual Review of Entomology 61:335–352.

McGraw, K. A. 2009. The Olympia Oyster, Ostrea lurida Carpenter 1864 Along The West Coast Of North America. Journal of Shellfish Research 28:5–10.

Miralles, L., E. Dopico, F. Devlo-Delva, and E. Garcia-Vazquez. 2016. Controlling populations of invasive pygmy mussel (Xenostrobus securis) through citizen science and environmental DNA. Marine Pollution Bulletin 110:127–132.

Novoa, A., R. Shackleton, S. Canavan, C. Cybèle, S. J. Davies, K. Dehnen-Schmutz, J. Fried, M. Gaertner, S. Geerts, C. L. Griffiths, H. Kaplan, S. Kumschick, D. C. Le Maitre, G. J. Measey, A. L. Nunes, D. M. Richardson, T. B. Robinson, J. Touza, and J. R. U. Wilson. 2018. A framework for engaging stakeholders on the management of alien species. Journal of Environmental Management 205:286–297.

Overdevest, C., C. H. Orr, and K. Stepenuck. 2004. Volunteer Stream Monitoring and Local Participation in Natural Resource Issues. Human Ecology Review 11:9.

Padilla-Ramírez, S., F. Díaz, A. D. Re, C. E. Galindo-Sanchez, A. L. Sanchez-Lizarraga, L.A. Nuñez-Moreno, D. Moreno-Sierra, K. Paschke, and C. Rosas. 2015. The effects of thermal acclimation on the behavior, thermal tolerance, and respiratory metabolism in a crab inhabiting a wide range of thermal habitats (Cancer antennarius Stimpson, 1856, the red shore crab). Marine and Freshwater Behaviour and Physiology 48:89–101.

Polson, M. P., and D. C. Zacherl. 2009. Geographic distribution and intertidal population status for the Olympia oyster, Ostrea lurida Carpenter 1864, from Alaska to Baja. Journal of Shellfish Research 28:69–77.

R Core Team. 2020. R: A language and environment for statistical computing. R Foundation for Statistical Computing, Vienna, Austria.

Roos, A. M. D., T. Schellekens, T. van Kooten, K. van de Wolfshaar, D. Claessen, and L. Persson. 2007. Food_JDependent Growth Leads to Overcompensation in Stage_JSpecific Biomass When Mortality Increases: The Influence of Maturation versus Reproduction Regulation. The American Naturalist 170:E59–E76.

Schneider, C. A., W. S. Rasband, and K. W. Eliceiri. 2012. NIH Image to ImageJ: 25 years of image analysis. Nature Methods 9:671–675.

Seebens, H., T. M. Blackburn, E. E. Dyer, P. Genovesi, P. E. Hulme, J. M. Jeschke, S. Pagad, P. Pyšek, M. Winter, M. Arianoutsou, S. Bacher, B. Blasius, G. Brundu, C. Capinha, L. Celesti-Grapow, W. Dawson, S. Dullinger, N. Fuentes, H. Jäger, J. Kartesz, M. Kenis, H. Kreft, I. Kühn, B. Lenzner, A. Liebhold, A. Mosena, D. Moser, M. Nishino, D. Pearman, J. Pergl, W. Rabitsch, J. Rojas-Sandoval, A. Roques, S. Rorke, S. Rossinelli, H. E. Roy, R. Scalera, S. Schindler, K. Štajerová, B. Tokarska-Guzik, M. van Kleunen, K. Walker, P. Weigelt, T. Yamanaka, and F. Essl. 2017. No saturation in the accumulation of alien species worldwide. Nature Communications 8:14435.

Simberloff, D. 2003. Eradication—preventing invasions at the outset. Weed Science 51:247– 253.

Simberloff, D., J.-L. Martin, P. Genovesi, V. Maris, D. A. Wardle, J. Aronson, F. Courchamp, B. Galil, E. García-Berthou, M. Pascal, P. Pyšek, R. Sousa, E. Tabacchi, and M. Vilà. 2013. Impacts of biological invasions: what’s what and the way forward. Trends in Ecology & Evolution 28:58–66.

Sorte, C. J. B., I. Ibanez, D. M. Blumenthal, N. A. Molinari, L. P. Miller, E. D. Grosholz, J. M. Diez, C. M. D’Antonio, J. D. Olden, S. J. Jones, and J. S. Dukes. 2013. Poised to prosper? A cross-system comparison of climate change effects on native and non-native species performance. Ecology Letters 16:261–270.

Sulkin, S. D., and G. McKeen. 1994. Influence of temperature on larval development of four co-occurring species of the brachyuran genus Cancer. Marine Biology 118:593–600.

Wasson, K. C, Zabin Bible, J., Ceballos, E., Chang, A., Cheng, B., Deck, A., Grosholz, E., Latta, M., and Ferner, M. 2014. A Guide to Olympia Oyster Restoration and Conservation.

Wasson, K., B. B. Hughes, J. S. Berriman, A. L. Chang, A. K. Deck, P. A. Dinnel, C. Endris, M. Espinoza, S. Dudas, M. C. Ferner, E. D. Grosholz, D. Kimbro, J. L. Ruesink, A. C. Trimble, D. V. Schaaf, C. J. Zabin, and D. C. Zacherl. 2016. Coast-wide recruitment dynamics of Olympia oysters reveal limited synchrony and multiple predictors of failure. Ecology 97:3503–3516.

Wetlands and Water Resources, Inc. 2010. Aramburu Island Shoreline Protection and Ecological Enhancement Project Draft Enhancement Plan.

Wickham, H., M. Averick, J. Bryan, W. Chang, L. D. McGowan, R. François, G. Grolemund, A. Hayes, L. Henry, J. Hester, M. Kuhn, T. L. Pedersen, E. Miller, S. M. Bache, K. Müller, J. Ooms, D. Robinson, D. P. Seidel, V. Spinu, K. Takahashi, D. Vaughan, C. Wilke, K. Woo, and H. Yutani. 2019. Welcome to the Tidyverse. Journal of Open Source Software 4:1686.

Wray, G. A., and R. A. Raff. 1991. The evolution of developmental strategy in marine invertebrates. Trends in Ecology & Evolution 6:45–50.

Zipkin, E. F. Z. F., P. J. S. J. Sullivan, E. G. C. G. Cooch, C. E. K. E. Kraft, B. J. S. J. Shuter, and B. C. W. C. Weidel. 2008. Overcompensatory response of a smallmouth bass (Micropterus dolomieu) population to harvest: release from competition? Canadian Journal of Fisheries and Aquatic Sciences 65:2279–2292.

Zu Ermgassen, P. S. E., M. D. Spalding, B. Blake, L. D. Coen, B. Dumbauld, S. Geiger, J. H. Grabowski, R. Grizzle, M. Luckenbach, K. McGraw, W. Rodney, J. L. Ruesink, S. P. Powers, and R. Brumbaugh. 2012. Historical ecology with real numbers: past and present extent and biomass of an imperilled estuarine habitat. Proceedings of the Royal Society B-Biological Sciences 279:3393–3400.

